# High-fidelity Image Restoration of Large 3D Electron Microscopy Volume

**DOI:** 10.1101/2023.09.14.557785

**Authors:** Yuri Kreinin, Pat Gunn, Dmitri Chklovskii, Jingpeng Wu

**Affiliations:** LDV Soft Inc, Aurora, Canada L4G6L6; Center for Computational Neuroscience, Flatiron Institute, New York, NY, USA 10010; Scientific Computing Core, Flatiron Institute, New York, NY, USA 10010; Neuroscience Institute, New York University School of Medicine, New York, NY 10016

**Author notes:** Electronic mail (Dmitri Chklovskii).

**Keywords:** Image Processing, Volume Electron Microscopy, Connectomics, Image Restoration

## Abstract

Volume Electron Microscopy (VEM) is an essential tool for studying biological structures. Due to the challenges of sample preparation and continuous volumetric imaging, image artifacts are almost inevitable. Such image artifacts complicate further processing both for automated computer vision methods and human experts. Unfortunately, the widely used Contrast Limited Adaptive Histogram Equalization (CLAHE) can alter the essential relative contrast information about some biological structures. We developed an image-processing pipeline to remove the artifacts and enhance the images without CLAHE. We apply our method to VEM datasets of a Microwasp head. We demonstrate that our method restores the images with high fidelity while preserving the original relative contrast. This pipeline is adaptable to other VEM datasets.

## Introduction

As biological structures are three-dimensional, Electron Microscopy technology has been extended for 3D volumetric imaging, called Volume Electron Microscopy (VEM) (Peddie *et al*., 2022; Peddie and Collinson, 2014; Chklovskii, Vitaladevuni, and Scheffer, 2010; Plaza, Scheffer, and Chklovskii, 2014). With the increase of both resolution and sample size, VEM is playing an increasingly important role in exploring biological structures and beyond.

A major motivation to develop VEM technology is connectomics, a field of neuroscience that aims at the dense reconstruction of neural networks in brains. Currently, large-scale VEM is still the only practical approach to acquiring connectomes (Peddie *et al*., 2022; Peddie and Collinson, 2014; Winding *et al*., 2022; Helmstaedter, 2013; Denk, Briggman, and Helmstaedter, 2012; DeFelipe, 2010; Plaza, Scheffer, and Chklovskii, 2014). Neurons are the basic functional units of brains, and they connect each other with synapses. A connectome is essential for the understanding of brain function and dysfunction (Galili, Jefferis, and Costa, 2022). Due to the variation in the scale of synapses and neurons, imaging for connectomics is best performed with both a wide field of 3D view and high resolution. The resulting image volumes are large-scale and challenging to analyze (Lichtman and Denk, 2011). Despite significant progress in sample preparation and imaging, image artifacts could still be found in image datasets due to various factors, including sample uniformity and imaging longevity. The artifacts could not only reduce the accuracy of automated detection and segmentation but also challenge human proofreaders.

Recently, we have acquired several 3D isotropic whole-brain datasets of Microwasp using customized VEM to construct connectomes (Makarova, Polilov, and Chklovskii, 2021). The tiny size of the Microwasp created immense challenges in sample preparation and whole-brain imaging, there exist several issues in the raw images (Fig. 1): (1) there exist strip artifacts; (2) there exists a great difference in the average or background intensity across the neighboring sections; (3) the background intensity in each section is not uniform; (4) there exists noise; (5) the contrast is low.

**FIG. 1.**
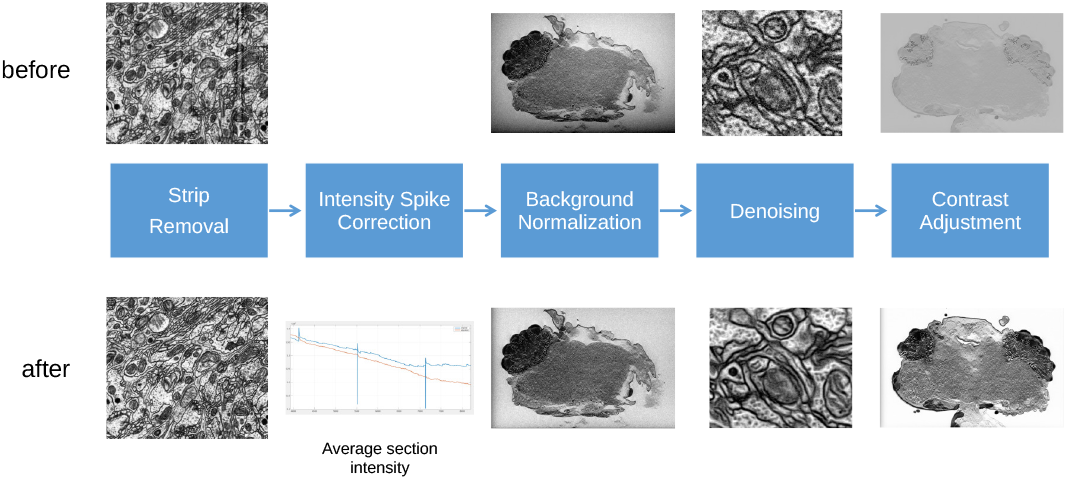
The whole pipeline of image processing. The effect of each step is illustrated by comparing the image and statistics before and after processing. The contrast of images is adjusted for visualization.

Interpolation is a priority in our image preprocessing pipeline to reduce generated artificial structures in the following biological studies. Recently developed Deep Neural Networks (DNNs) could be trained to restore images using either supervised (Fang *et al*., 2021) or self-supervised approaches (Buchholz *et al*., 2019). A Cycle Generative Adversarial Network (CycleGAN) could also be trained to transfer the image styles across different image datasets (Zhu *et al*., 2017; Scheffer *et al*., 2020). However, DNN training is still poorly understood, and model Interpretability is a well-known issue (Zhang and Zhu, 2018). There is a potential risk of hallucination when “imagined” structures could be generated, which could mislead biological discoveries. Despite the rapid development of image preprocessing techniques using DNN in the computer vision field, traditional methods remain the mainstay first approach in most large-scale connectomics projects (Macrina *et al*., 2021; and J. Alexander Bae *et al*., 2021; Shapson-Coe *et al*., 2021; Zheng *et al*., 2018; Scheffer *et al*., 2020; Chklovskii, Vitaladevuni, and Scheffer, 2010; Plaza, Scheffer, and Chklovskii, 2014). Thus, we initially attempt to preprocess data using non-DNN image processing approaches and only consider DNN when traditional approaches fail. Here, we have developed a customized image preprocessing pipeline to reduce five types of image artifacts.

## Materials and Methods

### A Whole Microwasp Brain Dataset

Microwasp, *Megaphragma vigianii*, is a compelling object of study for neuroscience (Makarova, Polilov, and Chklovskii, 2021) because of the small number (∼8,000) neurons in its brain, elaborate behavior, and miniaturization adaptations, such as the lack of most nuclei (Polilov, 2012, 2015). Using an ad hoc protocol the whole head of the Microwasp was prepared for EM by staining with a heavy metal (Polilov *et al*., 2021). The sample was imaged using a customized Focused Ion Beam Scanning Electron Microscopy (FIB-SEM) with a voxel size of 8 ×8 ×4 nm (Knott *et al*., 2008; Xu *et al*., 2017) and a bit depth of 16. The images were aligned using affine transformations based on Scale-Invariant Feature Transform features (Saalfeld *et al*., 2010; Lowe, 1999). The neighboring sections were combined by averaging resulting in an isotropic voxel size of 8 ×8 ×8 nm. The resulting total volume size is about 1.4 teravoxels.

### Image Processing Pipeline

As illustrated in Fig. 1, our image processing pipeline comprises five steps: Stripe Removal, Intensity Spike Correction, Background Normalization, Denoising, and Contrast Adjustment. Note that all the processing steps before the final Contrast Adjustment are conducted with a high bit depth to reduce information loss. We detail the algorithms as follows.

#### 1. Stripe Removal

Stripes in 2D images can be detected as high-frequency components in the Fourier transform of the images (Fig. 2A,B). The power spectrum can provide information about the spatial frequency content of the stripe artifact, including its location and shape. Typically, high-frequency components in the spectrum correspond to the spatial frequency of the stripes in the image, which is indicative of the stripe artifact. In Fig. 2A,B, the vertical stripes in the spectrum of an image appear as periodic fluctuations in the horizontal direction. These fluctuations indicate the presence of high-frequency components in the image that are oriented vertically, suggesting that there exists an artifact or noise with vertical feature orientation.

**FIG. 2.**
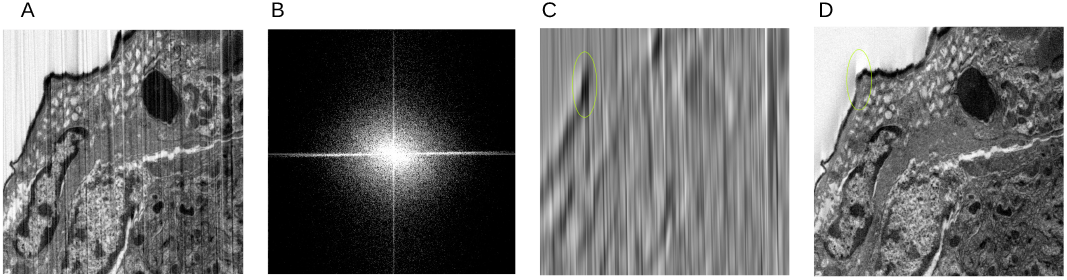
(A) Raw Image Impacted by Stripes. (B) The spectrum of A, (C) Stripe Pattern Extracted. The green circle indicates a membrane region with the same orientation of stripes and the signal was reduced after destriping. (D) De-striped Image obtained by subtraction of C from A.

Fig. 2A,B shows the strip artifact in our datasets in both spatial and frequency domains. One approach for reducing heavy stripes artifacts is to use image filtering algorithms such as Fourier filtering. These algorithms can remove high-frequency noise and smooth out variations in the image signal to reduce the stripes artifact, but they must be monitored against the possibility of introducing additional artifacts or distortions in the image. Instead of the widely used Nortch filter or Band-stop filter (Hirano, Nishimura, and Mitra, 1974), we designed a customized spatial domain model.

We assume that the distorted image is a composition of the stripe-free image and an additive symmetrical random noise term, that introduces the stripes.

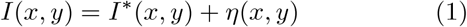

where *I* (*x, y*) is the observed pixel intensity value in location (*x, y*), *I*^∗^ (*x, y*) is original pixel value and *η* (*x, y*) is an additive noise, responsible for stripes artifact.

The fact that the stripes are primarily on the Y axis implies that the noise function *η* (*x, y*) has a much higher rate of change on the X axis than the Y axis. This leads to the following equation:

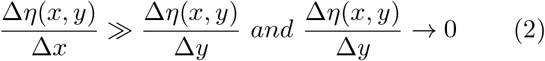

Equation 2 results in

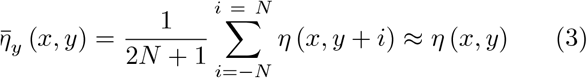

We also may assume that

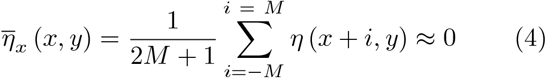

for *M* sufficiently large to satisfy the assumption that the noise is zero-mean.

The technique of using a moving average for equations 3 and 4 can be generalized by convolving the signal with low-pass filter *LP*_*y*_ and *LP*_*x*_. The equations can then be re-written as follows:

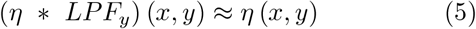

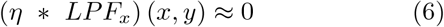

By using lowpass kernel *LPF* and its high pass complement *HPF*, which satisfies condition (*LPF* + *HPF*) (*x, y*) = 1, we can derive the formula for approximation of stripes noise from the observed signal (image). We rewrite equation 6:

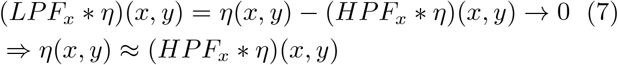

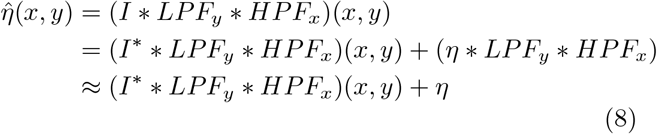

It appears that the spatial domain technique for estimating stripe noise aligns with the frequency domain analysis of the stripe noise spectrum, which was discussed earlier.

The equation 8 shows that stripe estimation is distorted by the term (*I*^∗^∗*LPF*_*y*_ ∗*HPF*_*x*_)(*x, y*). The convolution of image *I*^∗^(*x, y*) with the zero-phase low-pass filter *LPF*_*y*_ and high-pass filter *HPF*_*x*_ can result in the removal of useful information from the image, particularly in areas with strong vertical edges. This is due to the fact that the high-pass filter enhances high-frequency components in the x-direction, including stripes and vertical edges, while the low-pass filter smooths out these details in the vertical direction. While this smoothing process is effective at removing small and fine details from the stripe estimation, it does not affect long vertical edges. As a result, edge information is lost, which leads to a loss of image detail.

We employ a multi-scale decomposition approach to address the negative impact of vertical edges on the stripe detection process. This method involves analyzing the image at multiple scales, capturing larger-scale structures and finer details. We then adjust the filtering parameters accordingly using spatially adaptive filtering techniques. This allows for more precise targeting of the stripe artifacts in different image regions, yielding better performance compared to traditional fixed-filtering methods.

The decision to apply multi-scale decomposition is based on the assumption that stripes can be approximated by 1-dimensional white noise. The magnitude of this noise at each scale is determined by the frequency characteristics of the convolution kernels used for the decomposition, while the magnitude of the actual information in the signal depends on the image content. By utilizing multi-scale decomposition, we can effectively separate stripe artifacts from the image content and apply appropriate filtering to remove stripes without significantly affecting the underlying information.

Below is a brief summary of the steps implemented by the proposed solution:

1. The input image is first decomposed into multiple scales using a bank of separable filters. The separable filters can be used to create the next scale by convolving the previous scale with the the one-dimensional kernel in the y-direction and then convolving the result with the same kernel in the x-direction.
2. During this decomposition, the Initial Stripe Approximation Map is generated for each scale as the residual obtained by subtracting the next scale image from the intermediate result of convolution in the y-direction.
3. A threshold value is calculated for each row in the Initial Stripe Approximation Map.
4. The threshold values are used to convert the Initial Stripe Approximation Map into a binary map, with a value of 1 assigned to pixels that require caution during processing.
5. Connected components are extracted from the binary map and region properties such as orientation and major/minor axis length are calculated for each component.
6. The “stripness” confidence in the range of 0 to 1 is calculated for each connected component based on this region’s properties. The “stripness” confidence of 1 is assigned to the pixels that are marked as zero in a binary map.
7. The stripness confidence map is smoothed with a block filter and the Attenuated Stripe Approximation Map is generated by multiplying the Initial Stripe Approximation Map with the corresponding smoothed confidence map.
8. The Attenuated Stripe Approximation Map is smoothed with a vertical low-pass filter. We implement this filter by double-convolving the map with a vertical average kernel of length *M* ^(*n*)^, which is selected for each scale. The resulting image estimates the stripes artifact for the corresponding scale, which is then subtracted from the original image to suppress the stripes present in that particular frequency band.

**Multi-scale decomposition** can be performed using various techniques, including but not limited to Gaussian Pyramid (Burt, 1983), Laplacian Pyramid (Burt, 1983), and Wavelet Transform (Mallat, 1989).

We performed the multi-scale decomposition using a 7-tap low-pass filter (Fig. 3,4). This filter is specifically designed to avoid aliasing, making it well-suited for the atrous convolution technique. With this technique, we can efficiently compute a lower-scale representation by convolving the previous scale with the same kernel, but with a dilation factor of 2 (Fig. 5).

**FIG. 3.**
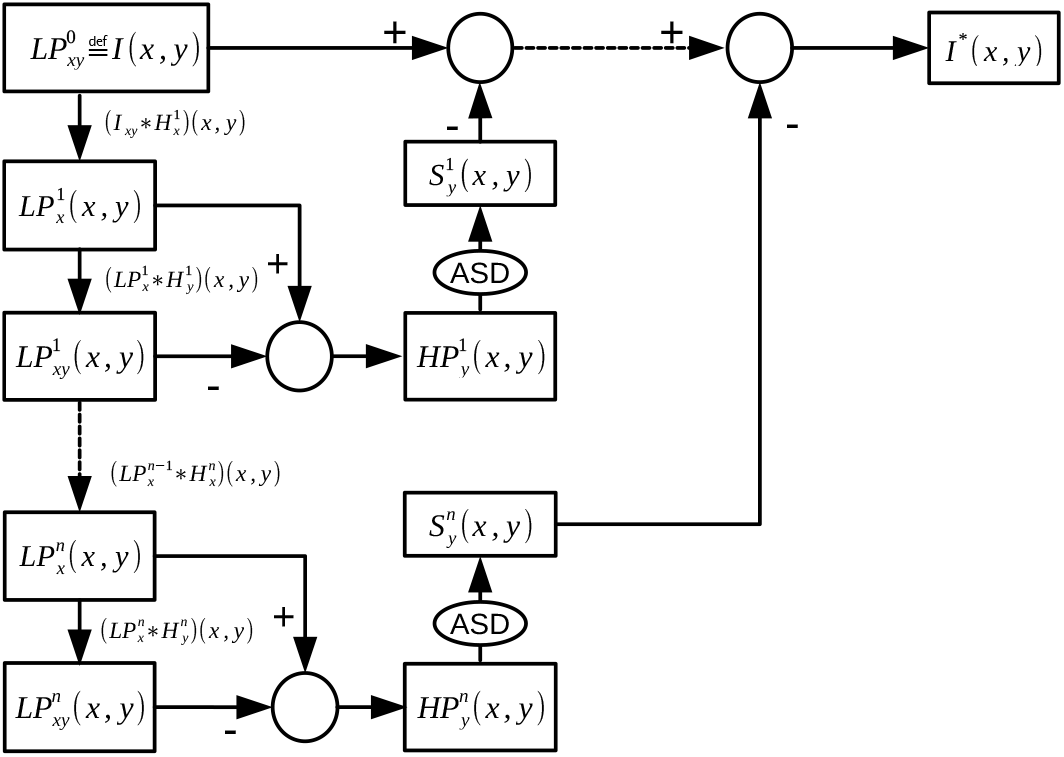
Multi-scale Decomposition for Stripe Artifact Suppression.

**FIG. 4.**
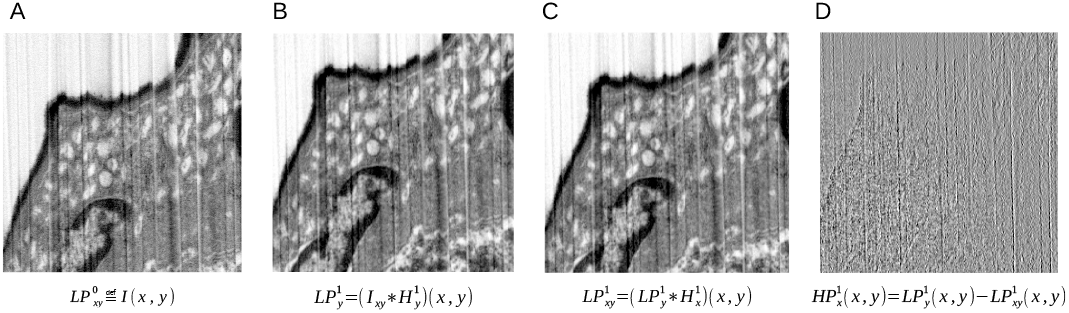
Sample images illustrating the decomposition steps at the 1st scale. (A) Raw image. (B) Lowpass filtered result in the Y direction. (C) Lowpass filtered result in the XY direction. (D) The difference between B and C.

**FIG. 5.**
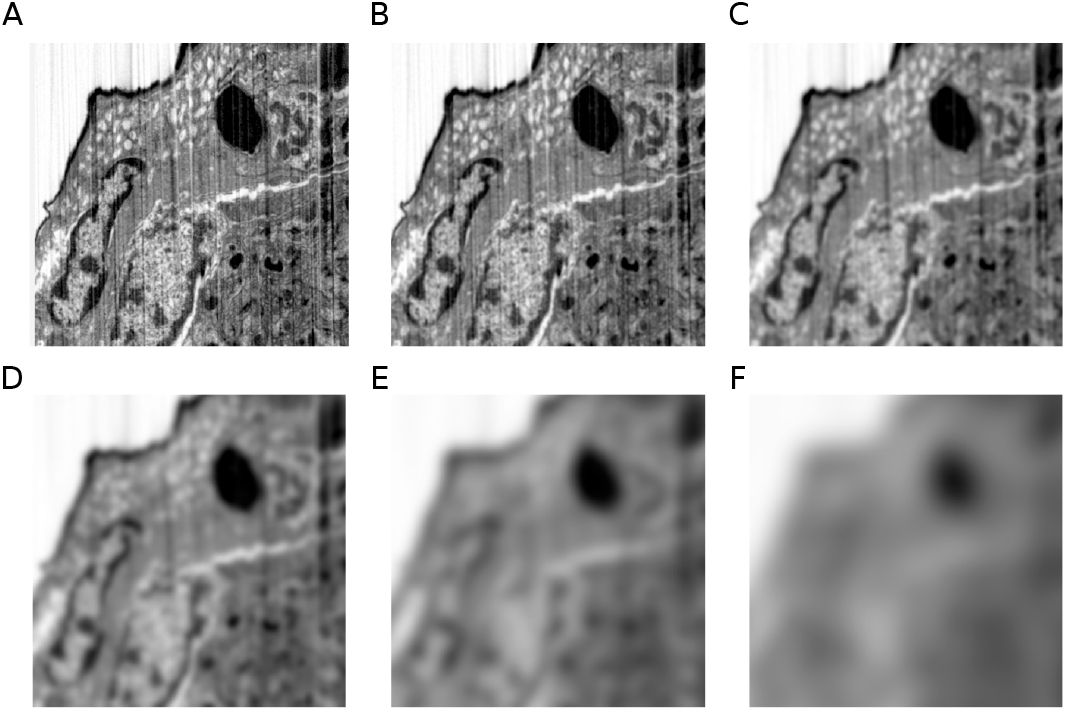
Low-pass filtered images 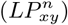 at different scales (n = 1 to 6 for A-F).

**The adaptive stripe detection** is performed independently for each scale. It converts 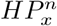 maps obtained for each scale into an accurate stripe approximation that mitigates the impact on vertical edges while preserving the actual stripes that need to be subtracted from the original image.

Based on our observations, the strength of the stripes is limited with respect to the signal and it changes gradually in the vertical direction, while the magnitude is constant in the horizontal direction. In order to mitigate the impact on the real signal, we aim to reduce the contribution of 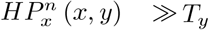, where 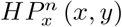 is the value of the Initial Stripe Approximation Map at location (x,y) for scale *n* in the multi-scale decomposition of the image (Fig. 6). *T*_*y*_ is threshold calculated for each horizontal row.

**FIG. 6.**
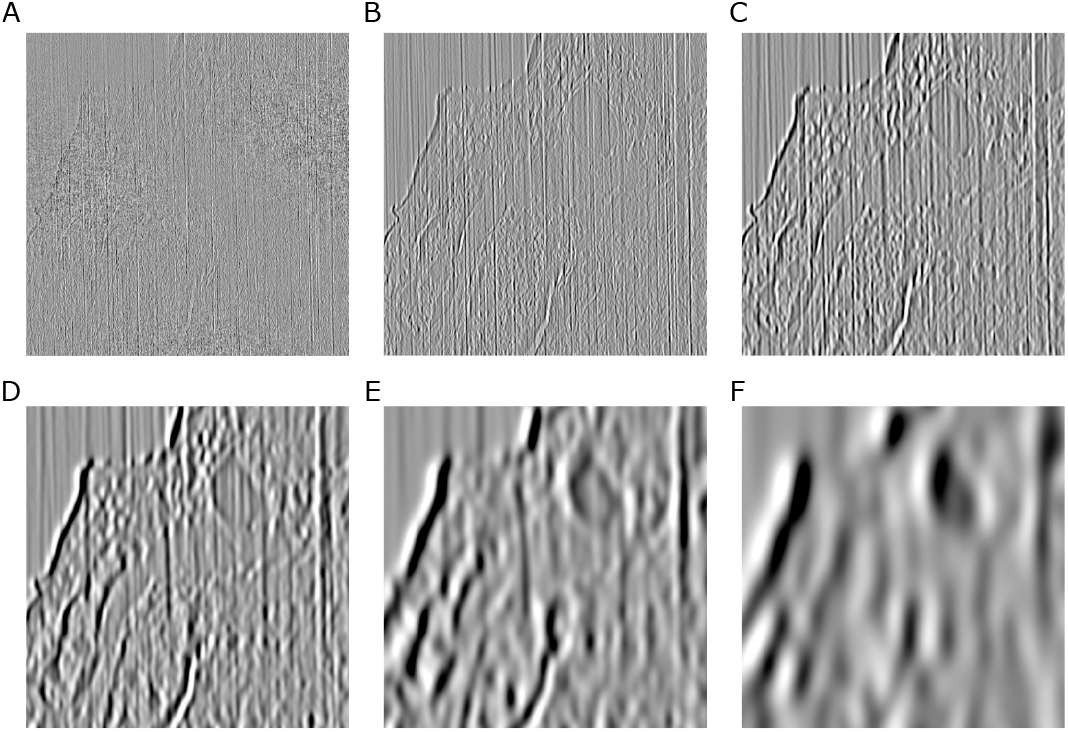
Stripe approximations 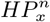 at various scales prior to further processing (n = 1 to 6 for A-F).

To obtain this adaptive threshold, we calculate the root mean square (RMS) of the values in the Initial Stripe Approximation Map for each row. The RMS is then smoothed using a moving average:

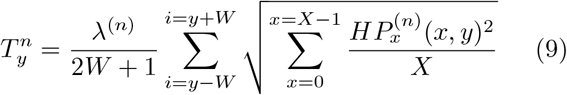

where: *X* is the number of columns (image width), *W* is an arbitrary number of rows before and after *y*, used to smooth the threshold. *λ* is a tuning parameter, heuristically selected for the scale *n*. The value for *λ* may vary in range from 1 to 3 depending on the observed strength of stripe artifacts.

The threshold value *T*_*y*_ is applied to create a binary map *BM* (*x, y*) from the initial stripe approximation map (Fig. 7). Each pixel in the binary map is set to 1 if the absolute value at the same location in the Initial Stripe Approximation Map is greater than the threshold value. This process generates a binary image where pixels with a value of 1 represent areas where caution is needed during processing. These areas may contain either stripes or edges, and their presence in the binary image indicates that this uncertainty needs to be resolved.

**FIG. 7.**
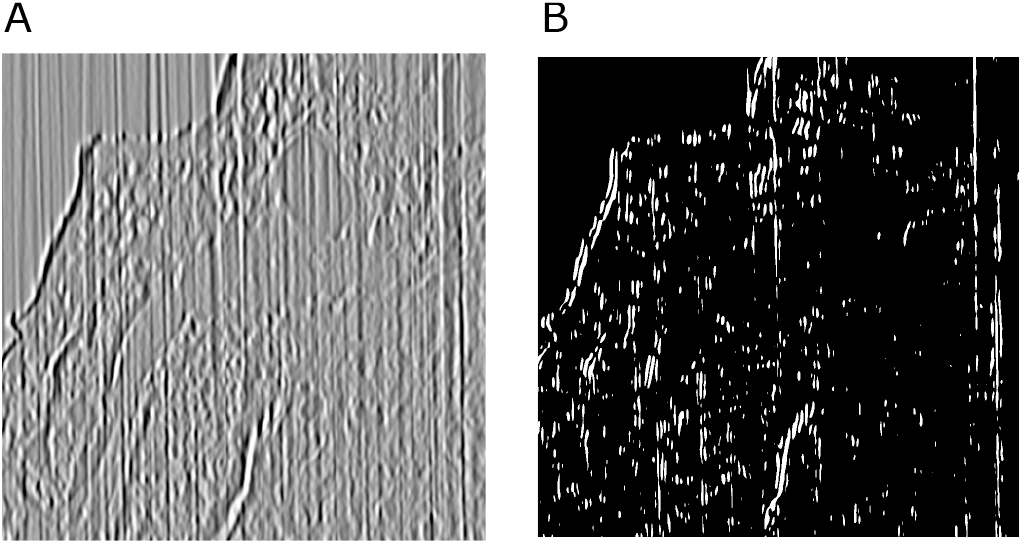
(A) Initial Stripes Estimation Map. (B) Binary map indicating uncertain pixels

The connected components (CC) are extracted from the binary map. For each connected component, we calculate a set of region properties consisting of *orientation, major*, and *minor axis length*.

- *Major axis length (a)* is length (in pixels) of the major axis of the ellipse that has the same normalized second central moments as the region.
- *Minor axis length (b)* is length (in pixels) of the minor axis of the ellipse that has the same normalized second central moments as the region.
- *Orientation (θ in* 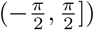) is the angle between the *x* -axis and the major axis of the ellipse that has the same second-moments as the region. It can be seen that connected components with an absolute orientation close to 90° are highly likely to be part of a stripe. The confidence decreases rapidly beyond a certain threshold, indicating that the connected component is not part of a stripe.
- *Aspect Ratio* 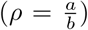, the ratio of the *major axis* to the *minor axis*. From our observations, it is apparent that a higher aspect ratio is indicative of a greater likelihood of the shape being a part of the stripe.

We assign stripness confidence values to connected components based on their region properties. Our observations suggest that connected components with a close-to-vertical orientation and a large aspect ratio are highly likely to be part of a stripe. However, the confidence rapidly drops after some deviation from a vertical orientation, and a decrease in aspect ratio. We also can preserve the contribution of connected components with a major axis significantly smaller than the length of the averaging filter (M), as their impact on the stripe estimation is mitigated by the averaging.

We introduce a heuristic that incorporates our observation. Two parameters are used to control the behavior of the output:

- *P* -minimal ratio between major and minor axes.
- Θ -minimal orientation angle

The confidence value for each connected component (*CC*_*i*_) is calculated, as:

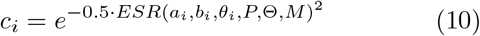

where *ESR* function measures the ratio between the presence of edges and the presence of stripes, which is the basis for the stripness confidence values associated with a connected component.

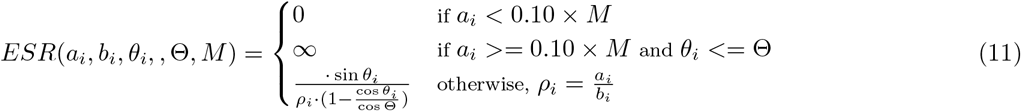

The fig. 8 illustrates curves of confidence value for different control parameters *P and* Θ.

**FIG. 8.**
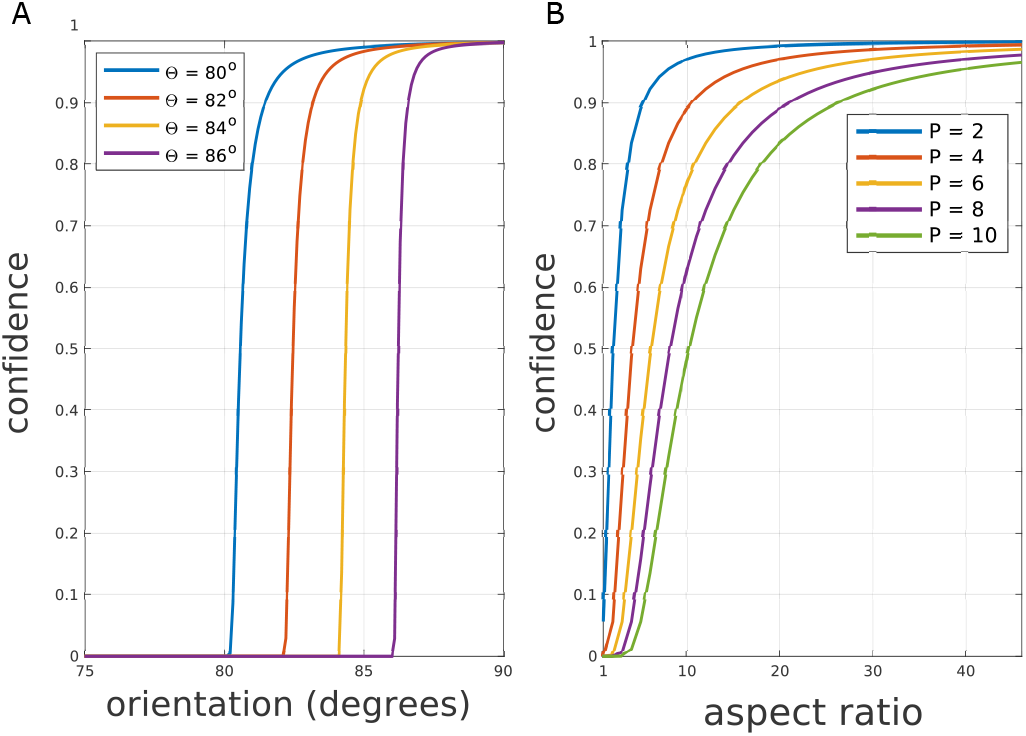
Relationship between confidence value and control parameters. (A) Change of confidence value for different minimal orientation (Θ), P = 10/3, ρ = 50/1. (B) change of confidence value for different minimal aspect ratios (P), θ = 89o,Θ = 84o

In order to mitigate the impact of the edge regions we create a map of weights *W* (*x, y*) for each location (x,y)

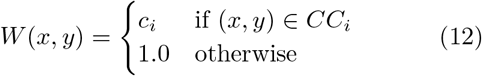

We smooth this weights map by convolving it with a rectangular block filter of size N × N, where N is selected for the best performance at each scale. The size of the filter increases with the order of scale to accommodate the increasing area of connected components. The Attenuated Stripe approximation Map (*ASM*) is generated by multiplying the Initial Stripe approximation Map 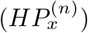 with the corresponding smoothed weights map.

Finally, the accurate stripe estimation for scale 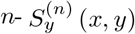 is obtained by smoothing the *ASM* with a vertical low-pass filter. This low-pass filter is implemented by two sequential convolution of *ASM* with vertical average kernel *A*^(*n*)^ of length *M* ^(*n*)^, where *M* is selected separately for each scale

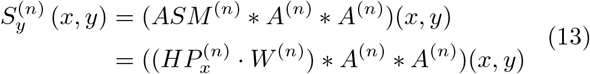

The resulting stripes map suppresses undesired impact of term (*I*^∗^ ∗*LPF*_*y*_ ∗*HPF*_*x*_)(*x, y*), introduced in equation 8.

#### 2. Intensity Spike Correction and Background Normalization

The FIB-SEM image stack exhibits intensity spikes between adjacent image sections and uneven illumination across a single section, which can be caused by various factors such as variations in imaging conditions or specimen preparation. These artifacts can negatively impact image analysis and segmentation. We apply appropriate image processing techniques to correct these artifacts and improve the overall quality of the image.

To correct the intensity spike artifact, we analyze the difference in mean intensity between each pair of neighboring sections. Small differences between sections that follow a gradual trend can be classified as not spike artifacts with a high degree of confidence. On the other hand, a large and abrupt difference indicates the presence of a spike artifact and should be replaced by the value that follows a natural trend of intensity change. The weight function 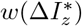 is calculated based on the magnitude of the difference in mean intensity between the two adjacent sections: 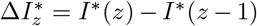. This weight function is defined such that it tends to be 1 for small values and drops to 0 for large ones. It reflects our assumption that the small difference in intensity it is likely a true change. The weight function is given by:

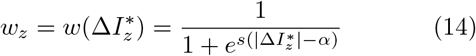

where *s* and *α* are parameters that control the steepness and sensitivity of the weight function to the magnitude of 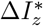.

By multiplying the mean intensity change 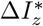 by the weight function, we obtain an adjusted value that estimates the confident change in intensity, meaning that we can be confident that this value is a reliable estimate of the true change:

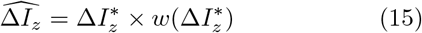

The final corrected intensity change between adjacent sections is calculated as (Fig. 9):

**FIG. 9.**
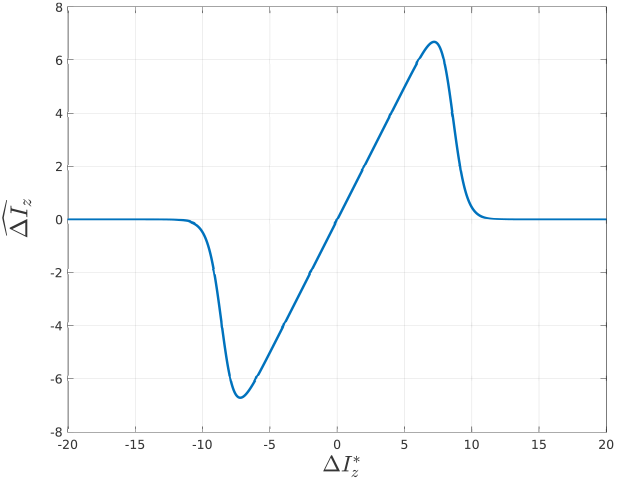
Estimation of the confident change in intensity (s = 2, α = 8.5)

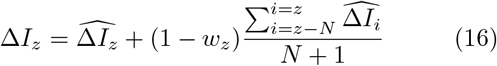

where *N* is parameter used to calculate the moving average of intensity differences for sections above z. The second term of the equation represents an estimation of the average change of the mean intensity of the N preceding sections, which accounts for any potential trend in the data.

### Background normalization

is the next step after spikes elimination. In order to perform background normalization, we use the binary mask of the wasp head created for segmentation purposes. Using this mask, we ignore the pixels belonging to the head and focus only on the background pixels. We then run a 3-dimensional polynomial fitting of the background pixels on the downsampled volume. This approach allowed us to achieve more accurate and efficient background normalization by avoiding interference from the foreground object (Fig. 10).

**FIG. 10.**
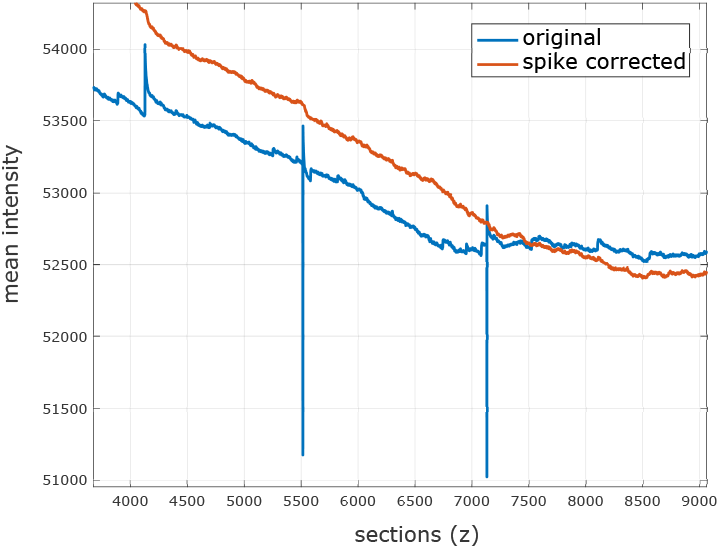
Mean intensity z-profile before and after spike correction.

#### 3. Denoising

The overall signal-to-noise ratio of the processed stack was reasonable, but to further reduce noise we applied a linear low-pass filter to suppress the highest frequencies, which are typically associated with noise. The filter was applied via a convolutional kernel in the spatial domain. Additionally, we performed a non-linear noise suppression step through local contrast enhancement, which will be discussed in the next section. These techniques allowed us to effectively reduce noise in the images while preserving important structures.

#### 4. Contrast Adjustment

The central goal of our connectomics is densely mapping the neurons and synapses. The mapping accuracy depends on the contrast of plasma membrane and T-bar (Svara *et al*., 2022; Lee *et al*., 2019). Thus, the contrast adjustment of the image should aim to improve the visibility of dark pixels while attenuating noise especially visible in bright areas. We performed this in two steps: Local Contrast Enhancement (LCE) and Global Contrast Enhancement (GCE).

**Local Contrast Enhancement** improves relative contrast between neighboring pixels. The LCE is applied on 2D images, i.e. for each section. The LCE method can be decomposed into the following steps:

1. The LCE image *I*_*LCE*_ (*x, y*) is calculated from the input image *I* (*x, y*) by applying the following formula:

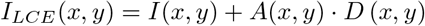

where *D* (*x, y*) are local details in location (*x, y*) and *A* (*x, y*) is an amplifier factor in the same location.
2. The details *D* (*x, y*) are calculated as a difference between pixel value *I* (*x, y*) and a middle value *M* (*x, y*) of the range in pixel’s neighborhood Ω.

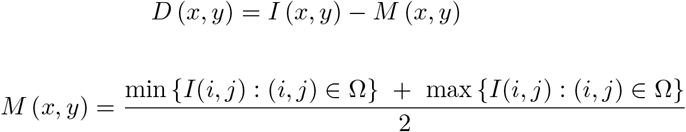

where set of pixel values *I* (*i, j*) is within the neighborhood Ω centered at pixel (*x, y*). For the actual dataset, the neighborhood Ω is 9 *×* 9 pixels.
3. In order to calculate non-linear amplifier factor *A* (*x, y*), we calculate the smoothed version *I*_*LP*_ (*x, y*) of the input image *I* (*x, y*) by applying a convolution with a low-pass gaussian filter *G* (*x, y, σ*_*LP*_): _*LP*_ (*x, y*) = (*I*∗ *G*) (*x, y*) The empirically chosen value of standard deviation *σ*_LP_ is 201 pixels.
4. The non-linear amplifier factor is computed by mapping the original pixel value *I* (*x, y*) to the output value *A* (*x, y*) using an S-shaped curve function. For the S-curve function, we picked the Complementary Cumulative Normal Distribution Function, also known as a survival function, which is the complement of the cumulative distribution function for the normal distribution *N* (*I*_*xy*_, *I*_*LP*_ (*x, y*) −*γ, σ*_*A*_), where *γ* and *σ*_*A*_ are parameters. This calculation is done by normalization of the input pixel value

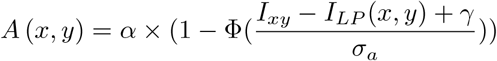

where Φ(*z*) is cumulative standard normal distribution function and *α, γ* and *σ*_*A*_ are parameters.

### Global Contrast Adjustment (GCA)

This approach not only improves the global contrast but also stretches the dynamic range of the image to the full available range from 0 to 255 since previous processing is performed with a high bit depth with a range of 0 to 1.0. The technique involves applying a transformation function to the pixel values of the image.

In order to enhance the contrast of an image while preserving its overall appearance and detail, the GCA transformation should satisfy the following criteria:

1. The transformation function should depend only on the pixel value: This means that the new pixel value in the output image is determined solely by the original pixel value in the input image, and not by the values of neighboring pixels or other factors. This ensures that the contrast adjustment is uniform across the image.
2. The transformation function used in GCA should be monotonically increasing: This means that as the pixel value increases in the input image, the corresponding mapped pixel value in the output image also increases. This is necessary in order to stretch the range of pixel values and enhance the contrast of the image, while preserving the relative intensities of the pixels.
3. The transformation function should preserve the relative intensities of the pixels in the input image: This means that the differences in intensity between adjacent pixels in the input image are maintained in the output image, even though the overall contrast of the image is enhanced. This is important for maintaining the visual quality of the image.
4. . The transformed pixel values should be within the valid range: The transformed pixel values should be within the valid range of [0, 255], which represents the range of possible pixel values in an 8-bit grayscale image.
5. The transformation function should be adjustable: The transformation function should be adjustable to allow for fine-tuning of the contrast adjustment based on the specific characteristics of the image being processed.
6. The transformation should avoid over-enhancement or artifacts, which can result in an unnatural or unrealistic-looking image.

Each pixel’s value is transformed using a linear equation that normalizes the pixel intensity to 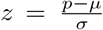, where p is the pixel intensity after applying Local Contrast Enhancement. The parameters *μ* and *σ* are determined based on the image histogram in the neuropil region. However, it’s important to note that these parameters are not necessarily the mean and standard deviation of the neuropil region itself, but rather they are used to normalize the pixel values across the entire image based on the statistics of the neuropil region.

After normalization is done, we apply a transformation function *H* (*z*) (Fig. 11), which is derived from the standard normal cumulative distribution function: 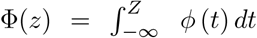 and standard normal probability distribution function 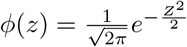

**FIG. 11.**
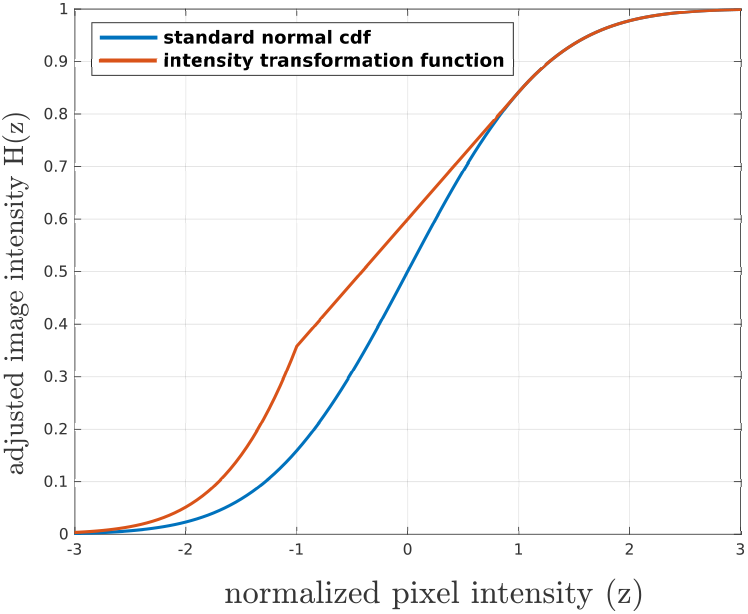
Transformation function H (z) (red) and standard normal cumulative distribution function (blue).

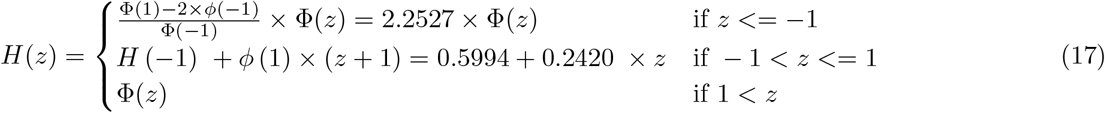

By using this transformation function, GCA is able to satisfy the criteria we discussed earlier, such as being a monotonically increasing function that depends only on the pixel value, preserving the relative intensities of the pixels, and being adjustable through the mean and standard deviation of the pixel values in the input image. This helps to bring out details and features that may be hidden in the original image due to low contrast.

The output of *H* (*z*) transformation is finally scaled to fit the range [0, 255] by multiplying it by 255.

## I Results

In order to illustrate the effectiveness of our method, we compared the results produced with Simple Global Contrast Adjustment (SGCA) and CLAHE. SGCA is a “levels” adjustment with the same parameters, applied to every image in the image stack. This was done by surveying a variety of brain areas and using the software GNU Image Manipulation Program (GIMP), an open-source image-editing tool similar to Adobe Photoshop, with its “curves” tool, to find upper, lower, and mean values to normalize the raw image data that result in data values suitable for humans to work with, and then applying those values to the full stack using the command line interface of ImageMagick. Since we have only adjusted the brightness and contrast, the SGCA result is close to the original raw images and is used as a reference for the metric of fidelity.

We compared the processing results of SGCA, CLAHE, and ours (Fig. 12). The section background intensity in the SGCA result is not even (Fig. 12 A) and is corrected by CLAHE (Fig. 12 B) and ours (Fig. 12 C). In the XZ and YZ sections of the SGCA result, there exists intensity variation across sections (Fig. 12 D, G), and is normalized by both CLAHE and our method. The relative darkness of some structures, such as eyes, are normalized by CLAHE, but conserved by our method (Fig. 12). The regions with less intensity variation become noisier after CLAHE processing.

**FIG. 12.**
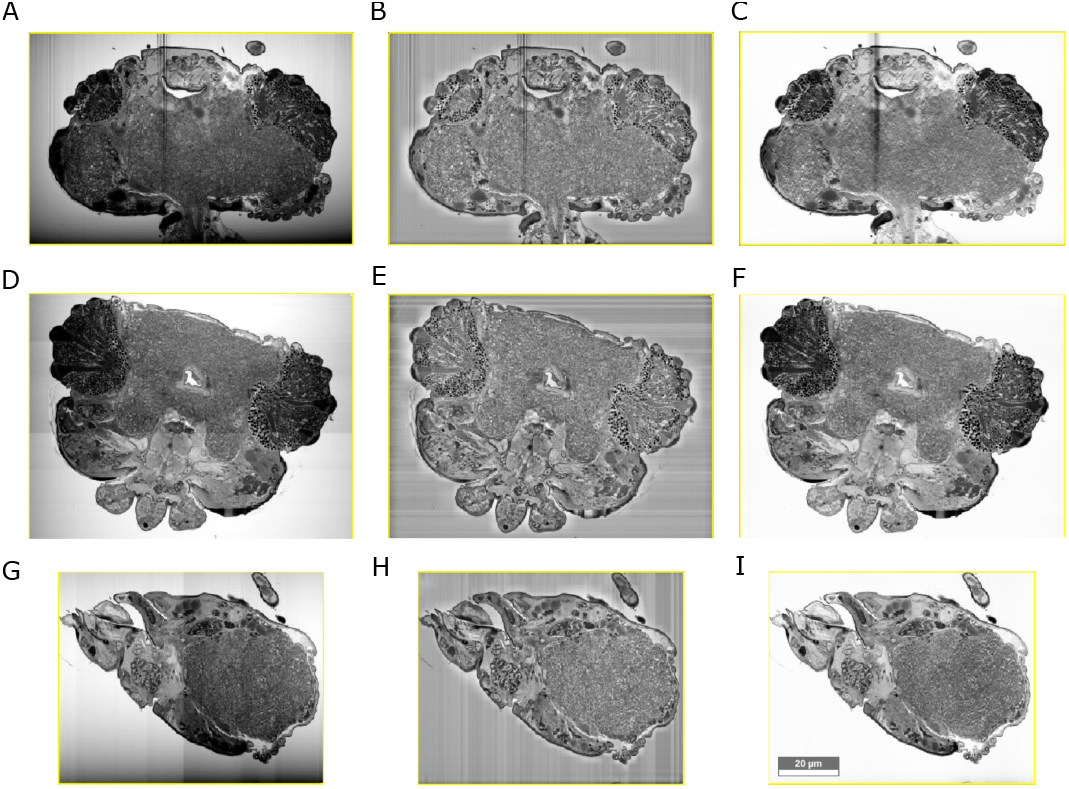
Cross sections of the whole Microwasp brain. From top to bottom, the sections are XY, XZ, and YZ planes respectively. From left to right, the sections are the results of SGCA, CLAHE, and our method respectively. The scale bar in I is 20 μm.

Despite the overview of whole head sections (Fig. 12), we also zoom in on some typical regions and compare the results (Fig. 13). In Fig. 13 A-C, the pigment cells and lens are consistently dark in SGCA and our result conserved this feature well with good fidelity. However, their relative intensity was changed by CLAHE with low fidelity. The lens and rhabdom were brightened by CLAHE processing as well. In Fig. 13 D-F, our stripe removal algorithm successfully restored the hidden image texture from the raw images with a high bit depth. In Fig. 13 G-I, CLAHE reduced the texture contrast but enhanced the noise of the background. Our method successfully enhanced the image texture contrast while reducing the noise level.

**FIG. 13.**
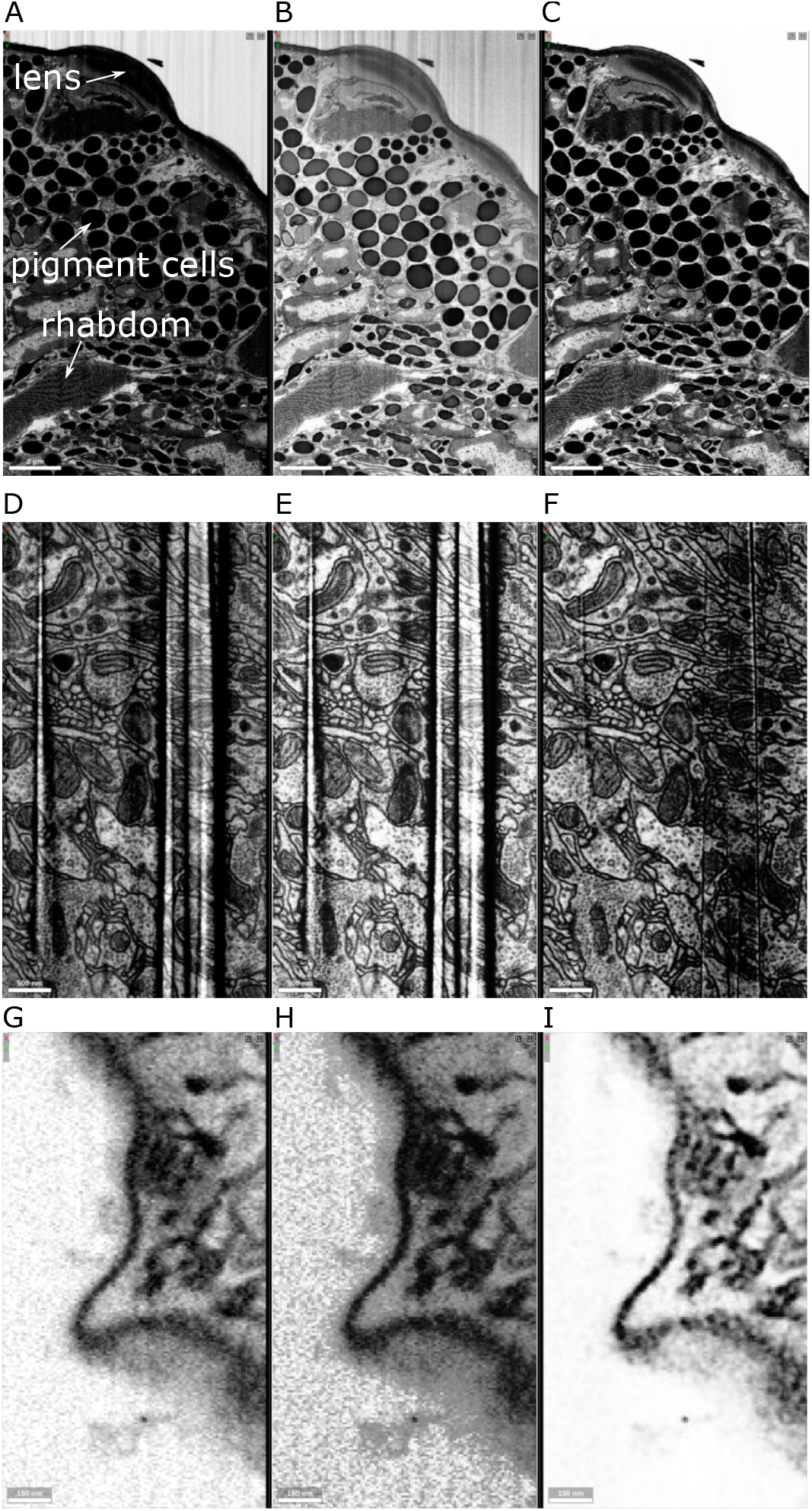
Region comparison of SGCA, CLAHE, and our method (from left to right). (A-C) An image region close to the eye. (D-F) An image region around the strips. (G-I) An image region covers the head boundary. From the top row to the bottom, the scale bars are 2 μm, 500 nm, and 150 nm respectively.

As illustrated in Fig. 14, the average intensity of sections along the Z-axis is uneven in the original images. With CLAHE, the average intensity is flattened and the intensity distribution becomes more uniform across the whole volume regardless nature of the biological structure (Fig. 14 B, D). For example, the darkness of cells in both eyes was normalized (Fig. 14 B). With our method, the average section intensity is normalized and the natural contrast of biological structures is preserved.

**FIG. 14.**
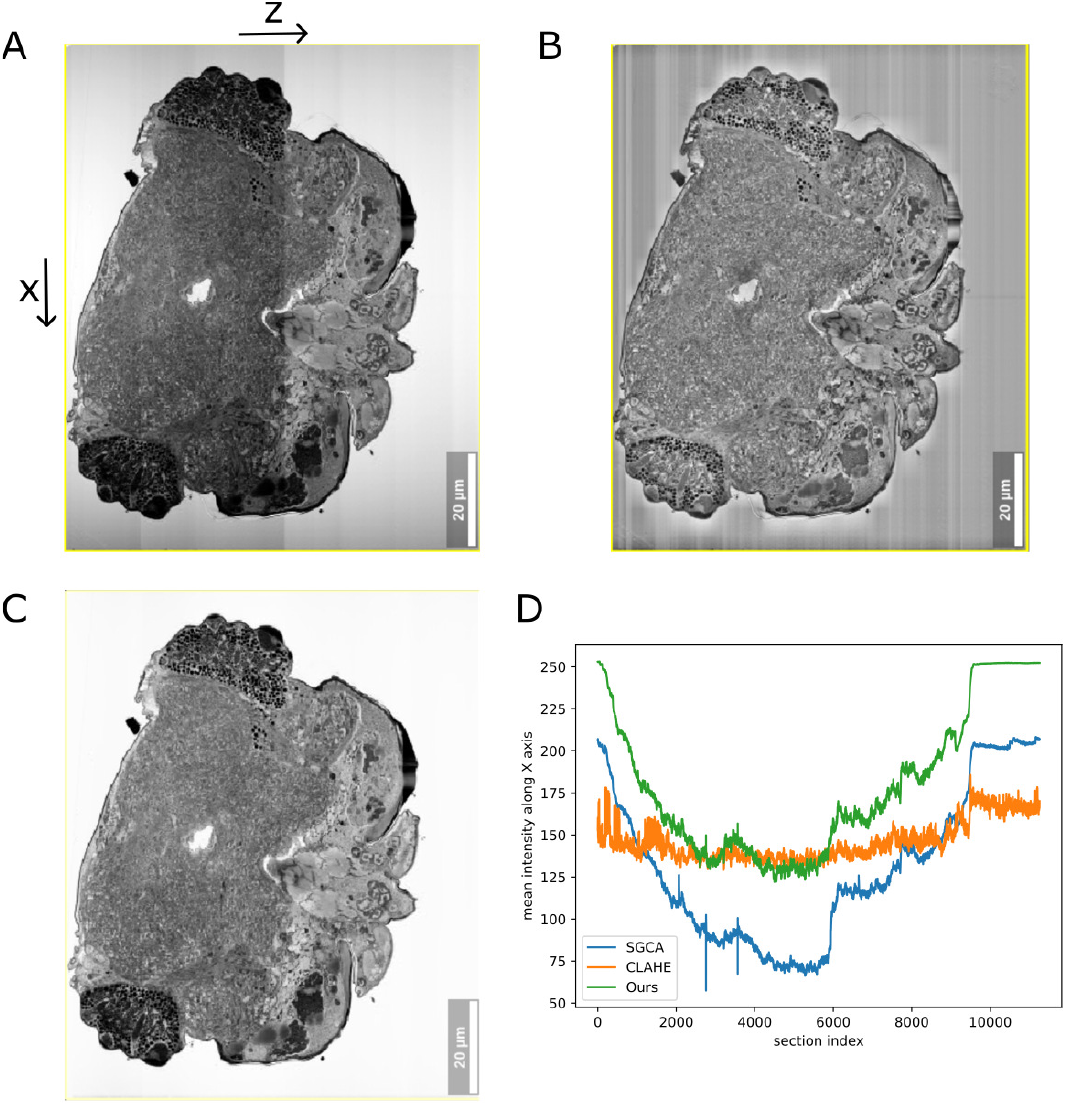
Comparison of average section intensity. (A-C) The XZ sections are in the center of the Y-axis of the whole dataset. (D) Comparison of average section intensity in A-C. The scale bars in A-C are 20 μm.

To address the data issues with our images, we developed an image processing pipeline that includes de-striping, intensity spike correction, background normalization, and contrast enhancement. We used this pipeline on our data and produced a high-quality image dataset with much fewer artifacts (Fig. 12, 13, 14).

## II Discussion

### II De-striping

There are various possible causes for stripe artifacts in FIB-SEM imaging. One such cause is uneven milling, leading to deep grooves or scratches on the sample surface. These irregularities can cause non-uniform removal of material during subsequent milling steps, resulting in heavy stripes in the final image. Another cause is sample surface charging during imaging, which can cause variations in the electron beam current and create non-uniform signals across the image. Other factors that can cause heavy stripes include improper SEM alignment, imaging parameter settings, and detector problems. Ways to minimize these artifacts include optimization of the FIB milling process, proper SEM alignment and tuning, and minimization of surface charging on sample surfaces. However, even with careful sample preparation and optimization of imaging parameters, the elimination of stripe artifacts is still a challenge.

Post-processing techniques such as image filtering or image stacking can be used to reduce or remove stripe artifacts from acquired images (Li *et al*., 2010; DING *et al*., 2013). These techniques can also smooth out variations in the image signal and improve the overall image quality.

Notch filtering, also known as band-stop filter, is a frequency-domain filtering technique that can be used to selectively remove or reduce specific frequency components from an image (Li *et al*., 2010; DING *et al*., 2013; Hirano, Nishimura, and Mitra, 1974). The technique starts with the creation of a filter with a narrow frequency band centered around the frequency of the artifact or noise that needs to be removed. To remove stripe artifacts, a notch filter can be designed to target the specific frequency of the stripes in the spectrum of the image. By removing the high-frequency components associated with stripes, the filter preserves the rest of the image content. A notch filter can also be applied in the spatial domain to achieve the same effect. Caution is necessary with this technique because over-filtering or inappropriate filter parameters can lead to the loss of vital image features or the introduction of additional artifacts. Bad design or application of the notch filter can adversely impact image quality. A common issue that can be caused by problems in notch filtering is the removal of critical image features or content that share the same frequency range as the artifact or noise being targeted. This can result in a loss of detail or distortion of the image, making it challenging to interpret or analyze.

A problem with the use of notch filters for stripe removal is that they are less effective at removing non-uniform stripes, which is a common issue in FIB-SEM imaging (Fig. 13D). Stripes introduced by FIB-SEM imaging are not evenly spaced and have varying amplitudes across the image.

To address this issue, we introduce an advanced filtering technique that is applied in the spatial domain with multiscale decomposition. Stripe Filtering in the spatial domain is not a new solution. However, we extend this technique by adding non-linear stripe suppression and by doing this in different frequency bands. This technique combines spatially adaptive filtering methods with the analysis of the image at multiple scales, enabling the adjustment of filtering parameters based on local statistical measures. By capturing both large-scale structure and fine details, this performs precise targeting of the stripe artifacts in different regions of the image. Compared to traditional fixed-filtering methods, this technique leads to improved performance in removing stripe artifacts.

Our method enables the analysis of an image across multiple scales, which facilitates the capture of both coarse-grained and fine-grained features. This multi-scale approach offers more precise targeting of stripe artifacts in various regions of the image, leading to better performance compared to conventional fixed-filtering techniques. The use of post-processing techniques can be an effective way to improve the quality of FIB-SEM images and reduce the impact of stripe artifacts.

### II Background normalization

The rolling ball algorithm is a popular method for background correction in microscopy images (Sternberg, 1983). However, it is best suited for images with a number of relatively small foreground objects, with similar areas and shapes. In the case of the wasp head electron microscopy image, object sizes vary across scales, examples being vesicles and cell bodies, both inside a single section and cross sections. Because of this, it is challenging to choose an appropriate size for structuring elements, and using a single structuring element may lead to incomplete or excessive background subtraction. To address this, we applied polynomial fitting.

### II Contrast adjustment

There are several off-the-shelf techniques that can be used for contrast enhancement, depending on the characteristics of the image and the specific features that need to be emphasized. Some of these techniques include histogram equalization, adaptive histogram equalization, contrast stretching, and gamma correction.

With our data, we found a combination of techniques to be the most effective. Local Contrast Enhancement is used to enhance the contrast of specific regions, such as those containing T-bars and membranes, while Global Contrast Adjustment via an S-shaped function is used to improve the overall contrast of the image. Parameters for these techniques should be optimized based on image characteristics and requirements of the segmentation task.

In many large-scale connectomics projects, a simple traditional method, Contrast-Limited Adaptive Histogram Equalization (CLAHE), has become the default image processing method due to its simplicity and computational efficiency (Pizer *et al*., 1987; Peddie *et al*., 2022; Scheffer *et al*., 2020). However, we found several issues while applying it to our dataset. First, the computation is normally performed tile by tile in large volumes. It normalizes a tile with close mean intensity regardless of the structure. If there are large dark structures with sizes comparable to the tile size, the relative intensity might be changed without respecting the physical property (Fig. 13 B). Second, the noise of the background or large objects could be amplified due to the dynamic local computation (Fig. 13 H). The parameter of histogram clipping may be based on the Probability Density Function or the percentage of the peak intensity. Fourth, the contrast is normalized completely locally and the contrast becomes even across the whole big volume which might not be consistent with the original biological structure. In our dataset, the biological (desired) intensity of the compound eye structures is much lower than neurites, while CLAHE-adjusted data did not retain this pattern (Fig. 12-14). Last, the contrast can be reduced rather than enhanced in areas where a large bright object, such as the background, is surrounded (Fig. 13 G-I). As illustrated in the results, our combined method avoided the above issues of CLAHE (Fig. 12-14).

### II Image processing with a high bit depth prevents information loss

In most EM connectomics projects, image processing was performed using gray-scale images with a bit depth of 8 (Turner *et al*., 2022; Macrina *et al*., 2021; and J. Alexander Bae *et al*., 2021). The bit depth of raw images produced by the microscope is normally higher with a much larger dynamic range. The information in such a high dynamic range will inevitably be lost during the quantization process. Thus, we perform all the image processing using a bit depth of 16 and quantize the image to 8-bit depth in the last step. Some image textures embedded in strips are successfully recovered using this approach (Fig. 13 D-F).

### II Limitations and remaining issues

Given a VEM dataset, we aim to restore the image signals as the first processing step. In contrast to Deep Learning approaches, we prefer traditional image-processing techniques due to their interpretability and flexibility of parameter adjustment. We would like to use traditional methods to solve as many issues as possible in this step. There still exist several issues and we plan to use Deep Learning approaches to handle them in the future.

In this initial step, our pipeline is designed only to restore images with high fidelity rather than inferring missing structures or warping the structures. As a result, the artifacts with morphological distortion are not handled. One such distortion is milling thickness variation. During EM imaging, a focused ion beam mills the surface of the sample, and the milling depth is not guaranteed to be consistent across either regions or sections. As a result, the surface is not guaranteed to be flat and the structures in regions of XZ and YZ sections are distorted. Since the thickness variation is inconsistent across regions and sections, a previous method is unsuitable for this dataset (Hanslovsky, Bogovic, and Saalfeld, 2016). Luckily, this artifact only exists in some small areas and most neurons are still traceable manually. Another issue is tissue damage. During sampling preparation, a small portion of brain tissue was damaged and the plasma membrane was broken. As both of these kinds of morphological distortion exist in some regions of our dataset, but don’t affect structure detection of most regions, we only plan to deal with them in the future if necessary.

### II Potential applications in other datasets

Although our method is designed to handle a specific dataset, it is still useful for other datasets with similar artifacts. The processing steps are relatively independent of each other and could be applied selectively. Because we apply these methods with a higher bit depth to reduce information loss, It is recommended to apply contrast enhancement and quantization in the final stage.

## III Conclusion

In order to remove artifacts in a whole Microwasp head dataset, we have developed an image processing pipeline. Compared with SGCE and CLAHE, our method is effective to remove the targeted artifacts and enhance the contrast respecting the original texture with high fidelity. The processing components are independent of each other and could be adopted selectively to other image datasets with similar artifacts.

## IV Code and Data Availability

The code, including a tutorial to process an image section, is available in a GitHub repository: https://github.com/brain-map/hifiem. The image section could be downloaded using the example Python Notebook. The whole brain datasets will be available upon the publication of another paper.

## V Acknowledgments

The sample preparation is supported by Russian Science Foundation (project no. 22-14-00028 to AP). The images were acquired and aligned by Song Pang and C. Shan Xu in the lab of Harald Hess at the Janelia Research Campus, which is supported by the Howard Hughes Medical Institute. The research at the Flatiron Institute is internally funded by the Simons Foundation.

